# *In vitro* evolution of ciprofloxacin resistance in *Neisseria* commensals and derived mutation population dynamics in natural *Neisseria* populations

**DOI:** 10.1101/2024.07.16.603762

**Authors:** Leah R. Robinson, Caroline J. McDevitt, Molly R. Regan, Sophie L. Quail, Crista B. Wadsworth

## Abstract

Commensal *Neisseria* are members of a healthy human oropharyngeal microbiome; however, they also serve as a reservoir of antimicrobial resistance for their pathogenic relatives. Despite their known importance as sources of novel genetic variation for pathogens, we still do not understand the full suite of resistance mutations commensal species can harbor. Here, we use *in vitro* selection to assess the mutations that emerge in response to ciprofloxacin selection in commensal *Neisseria* by passaging 4 replicates of 4 different species in the presence of a selective antibiotic gradient for 20 days; then categorized derived mutations with whole genome sequencing. 10/16 selected cells lines across the 4 species evolved ciprofloxacin resistance (≥ 1 ug/ml); with resistance-contributing mutations primarily emerging in *DNA gyrase subunit A* and *B* (*gyrA* and *gyrB*), *topoisomerase IV subunits C* and *E* (*parC* and *parE*), and the *multiple transferable efflux pump repressor* (*mtrR*). Of note, these derived mutations appeared in the same loci responsible for ciprofloxacin reduced susceptibility in the pathogenic *Neisseria*, suggesting conserved mechanisms of resistance across the genus. Additionally, we tested for zoliflodacin cross-resistance in evolved strain lines and found 6 lineages with elevated zoliflodacin minimum inhibitory concentrations. Finally, to interrogate the likelihood of experimentally derived mutations emerging and contributing to resistance in natural *Neisseria*, we used a population-based approach and identified GyrA 91I as a substitution circulating within commensal *Neisseria* populations and ParC 85C in a single gonococcal isolate. Small clusters of gonococcal isolates had commensal-like alleles at *parC* and *parE*, indicating recent cross-species recombination events.

## Introduction

Commensal species within the genus *Neisseria* have an intertwined relationship with their pathogenic relatives *Neisseria gonorrhoeae* (Ngo, the pathogen responsible for the sexually transmitted infection (STI) gonorrhea) and *N. meningitidis* (Nmen, the primary *Neisseria* species responsible for meningococcemia and bacterial meningitis), through widespread horizontal gene transfer (HGT) of both chromosomal^1–5^ and plasmid DNA^6^. Genetic exchange across species’ boundaries has led to the acquisition of important phenotypic traits, such as antibiotic resistance^1–5^, however, the commensal *Neisseria* are relatively underexplored for the phenotypic and genotypic diversity they harbor. Making up the most abundant genera within proteobacteria in oral and pharyngeal samples (∼10% of operational taxonomic units [OTUs]) in 100% of healthy human adults and children^7–11^, commensal species are a large portion of our oropharyngeal microbiomes; and thus will be a persistent threat for rapid pathogen evolution through bystander selection and DNA donation.

Though attention on the acquisition of antibiotic resistance through HGT between the *Neisseria* has often focused on beta-lactam resistance acquisition through mosaic *penicillin-binding protein 2* (*penA*)^4,12–15^ and macrolide resistance through inheritance of mosaic *multiple transferable efflux pump* (*mtr*) alleles^1,3,14,16,17^; quinolone resistance has also been documented to be transferred through transformation and homologous recombination^5^. Ciprofloxacin is a quinolone class antibiotic, which though once recommended as a first-line therapy to treat gonorrhea from the mid-late-1980’s, was discontinued in the 1990’s due to widespread resistance emergence^18^. The first clinical failures were reported by 1990 to low dose treatment (250 mg)^19^, which prompted a higher dose recommendation (500 mg), however resistance to high-dose treatment evolved rapidly in Asia^20^. The rate of ciprofloxacin resistance ≥ 1 ug/ml still remains high, with over 31% of U.S. isolates surveyed in 2022 classified as resistant^21^. Ciprofloxacin and other quinolone antibiotics target bacterial type II topoisomerases, DNA gyrase and topoisomerase IV, which disrupts DNA synthesis^22^. Work in *N. gonorrhoeae* has demonstrated that DNA gyrase subunit A (encoded by *gyrA*) is the primary target and topoisomerase IV subunit C (encoded by *parC)* is the secondary target for fluoroquinolones^23–25^. Unsurprisingly, reduced susceptibility to ciprofloxacin in gonococci tends to be conserved and emerge in the quinolone resistance-determining regions (QRDRs; codons 67–106 in GyrA and 56–108 in ParC) of the drug targets which reduce fluoroquinolone binding affinity^26–28^. Particular amino acid positions associated with reduced susceptibility include 91 and 95 in GyrA; and 85, 86, 87, 88, 91, and 116 in ParC; with genomic haplotypes harboring both *gyrA* and *parC* mutations displaying higher MICs than those with a single mutation^29^. Alternative pathways to ciprofloxacin reduced susceptibility exist however, with recent work in gonococci highlighting the emergence of mutations in DNA gyrase subunit B (GyrB D429N and P739H) and topoisomerase IV subunit E (ParE P456S) during *in vitro* selection^30,31^.

Acquisition of ciprofloxacin reduced susceptibility in both pathogenic *Neisseria* species has been associated with the inheritance of alleles from the commensals. For example, recent spread of commensally-acquired *gyrA* and *parC* haplotypes in meningococcal populations has played a major role in the establishment of quinolone resistant meningococci (QRM) globally^5,32,33^. The first records of commensally acquired quinolone resistance in meningococci are from the United States in 2007 and 2008. Three cases of meningococci from serogroup B were identified as ciprofloxacin-resistant (with MICs of 0.25 μg/mL) in two states (North Dakota and Minnesota)^34^. Currently the percentage of QRM with mosaic alleles is high, with over 53.4% and >70% of resistant isolates in the U.S. and China harboring mosaic alleles at *gyrA* and *parC*^5^. Furthermore, a global *in silico* analysis of >20,000 *Neisseria* isolates demonstrated that alleles at *gyrA*, *gyrB*, and *parE* have been transferred from commensal *Neisseria* to *N. gonorrhoeae*^35^. These examples underscore the importance of commensals in transferring quinolone resistance, and stress the importance of characterizing the quinolone-resistance conferring mutations that commensals may harbor.

Given the documented cross-species spread of quinolone resistance in natural *Neisseria* populations^5,35^, we set out to characterize the mutations which impart ciprofloxacin resistance in *Neisseria* commensals using *in vitro* selection. Here, we passage four species of commensals using a selective gradient of ciprofloxacin for 20 days, and characterize derived mutations post-selection. As in our prior work^36,37^, we query the identity and number of paths to reduced susceptibility, and compare and contrast these paths across different commensal genomic backgrounds. We also test for the potential for the emergence of zoliflodacin cross-resistance. Zoliflodacin is a spiropyrimidinetrione antibiotic which also binds type II topoisomerases, and has recently completed phase III trials for treatment of gonorrhea^38–40^. Though the mode of inhibition and binding sites are distinct from those of the fluoroquinolones^41^, we hypothesized that i) mutations impacting topoisomerase conformation or ii) general influx or efflux related mutations may emerge that could result in ciprofloxacin producing cross-resistance. Finally, we interrogate the presence of selected mutations in natural populations of *Neisseria*, and search for evidence of HGT from commensals to gonococci.

## Methods

### Bacterial Isolates

In this study, the following bacterial strains were examined; AR-0944 (*N. cinerea*), AR-0945 (*N. elongata*), AR-0948 (*N. canis*), and AR-0953 (*N. subflava*), which were acquired from the Centers for Disease Control and Prevention (CDC) and Food and Drug Association’s (FDA) Antibiotic Resistance (AR) Isolate Bank “*Neisseria* species MALDI-TOF Verification panel”. These strains were cultured on GC agar base (Becton Dickinson Co., Franklin Lakes, NJ, USA) media plates along with 1% BBL IsoVitaleX Enrichment (Becton Dickinson Co.,Franklin Lakes, NJ, USA; hereafter referred to as GCB-I media). These strains were grown for 18–24 hours at 37°C in a 5% CO_2_ incubator. Ancestral strain MIC values to ciprofloxacin were previously reported^42^ (Table 1), and confirmed prior to *in vitro* selection experiments. Stocks were stored at -80°C in a solution containing trypticase soy broth (TSB) and 20% glycerol.

**Table 1.** Characterization of ciprofloxacin MICs for ciprofloxacin-selected lineages.

|  | Ciprofloxacin MICs (µg/mL) |  |  |  |  |
| --- | --- | --- | --- | --- | --- |
| Species | Ancestral | Replicate 1 | Replicate 2 | Replicate 3 | Replicate 4 |
| AR-0944<br>(*N. cinerea*) | 0.032 | 4 | 12 | 8 | 6 |
| AR-0945<br>(*N. elongata*) | 0.25 | 0.125 | 2 | 1.5 | 0.75 |
| AR-0948<br>(*N. canis*) | 0.008 | 1.5 | 0.19 | 0.032 | 0.19 |
| AR-0953<br>(*N. subflava*) | 0.75 | 16 | 8 | 8 | 16 |

### In vitro selection of resistance to ciprofloxacin and antimicrobial susceptibility testing

The selection for ciprofloxacin resistance was performed via passaging four replicates of each of the aforementioned commensal strains in the presence of an ETEST strip to introduce a selective antibiotic gradient. In short, cells on the interior and from a 1 cm perimeter on the zone of inhibition (ZOI) were collected and suspended in TSB. This solution was then spread onto a fresh GCB-I plate, and incubated at 37°C with 5% CO_2_. 48 hours later, minimum inhibitory concentrations (MICs) were observed and recorded. The repetition of this procedure occurred for 11 passages over 21 days for each replicate. Bacterial populations from each replicate were passaged in the absence of antibiotic and struck out to collect single colonies, which were stocked for future analysis. Control strains were also passaged in the absence of ciprofloxacin with the same procedure.

The MIC values for the aforementioned post-selection individual colony stocks were measured with ETEST strips, which have generated comparable MIC results to the agar dilution method^43^. Simply put, bacterial cells from plates incubated overnight were suspended in TSB to a 0.5 McFarland standard, and then spread onto a GCB-I plate along with an Etest strip. After an 18–24 hour incubation period in the aforementioned conditions, MIC values were recorded and reduced susceptibility was determined using Clinical & Laboratory Standards Institute (CLSI) guidelines for *N*. *gonorrhoeae* of ≥ 1 μg/mL^44^. The final MIC values reported are the mode of three independent MIC tests and commonly agreed upon by two independent researchers.

### Genome sequencing and bioinformatics analysis for selected isolates

Bacterial cells from overnight plates were lysed via suspension in TE buffer (10 mM Tris [pH 8.0], 10 mM EDTA) with 0.5 mg/mL lysozyme and 3 mg/mL proteinase K (Sigma-Aldrich Corp., St. Louis, MO). DNA isolation was performed with PureLink Genomic DNA Mini Kit (Thermo Fisher Corp., Waltham, MA) with RNase A treatment for the removal of unwanted RNA. Isolated DNA was organized for subsequent sequencing utilizing the Nextera XT kit (Illumina Corp., San Diego, CA), and uniquely dual-indexed and pooled. The finalized pool was then sequenced using the Illumina MiSeq platform at the Rochester Institute of Technology Genomics Core using V3 600 cycle cartridges (2x300bp).

The quality of each paired-end read library was analyzed using FastQC v0.11.9. Any adaptor sequences or low-quality sequences were trimmed from read libraries using Trimmomatic v0.39, using a phred quality score < 15 over a 4 bp sliding window as a cutoff. Reads shorter than 36 bp, and those missing a mate, were removed from subsequent analysis as well. The reference genomes were assembled and annotated previously^36,37,42^, and the read libraries were mapped back to these reference assemblies using Bowtie2 v.2.2.4 using the “end-to-end” and “very-sensitive” options^45^. Ultimately, Pilon v.1.16^46^ identified any derived mutations; and custom scripts were used to determine if these mutations were located in any previously annotated genomic features. A common amino acid residue number scheme across species was determined by alignment to the NZ_AP023069.1 gonococcal reference genome.

### Testing for zoliflodacin cross-resistance

The presence of cross-resistance to zoliflodacin in the ciprofloxacin-selected replicates was determined experimentally using agar dilution. In brief, GC agar base with 1% IsoVitaleX was supplemented with zoliflodacin to produce plates containing 0.0, 0.06, 0.125, 0.25, 0.5, 1.0, 2.0, 4.0, and 8.0 µg/mL of drug. Bacterial cells from overnight plates were collected and suspended in 1 mL TSB and spotted on plates. Plates were incubated at 37°C with 5% CO_2_ for 16 hours, at which time MICs were recorded. Three independent MIC readings were recorded for each strain, with the mode reported as the final MIC. While the resistance breakpoint for species of *Neisseria* in the presence of zoliflodacin has not yet been defined, we use a prior published breakpoint of MIC of ≥ 4 µg/mL to define resistance^47^.

### Comparative population analysis of the presence of derived mutations and HGT in natural Neisseria populations

Previous work by Manoharan-Basil *et al.* (2022) has demonstrated alleles of *gyrA*, *gyrB*, and *parE* have been transferred from commensal *Neisseria* to *N. gonorrhoeae* in a population dataset from the 1920’s to 2020^48^. Here, we reevaluate cross-species sharing of these and *parC* alleles using isolates deposited to the PubMLST database (https://www.pubmlst.org/Neisseria)^49^, but focusing on the more recent depositions from 2020 through 2024. We specifically interrogate the presence of experimentally derived mutations in this dataset from our *in vitro* selection approach, and also search for any evidence of HGT of these particular alleles if they were present. 2235 *gyrA* (NEIS1320), 2302 *gyrB* (NEIS0204), 2287 *parE* (NEIS1600), and 2256 *parC* (NEIS1525) alleles were downloaded from published genomes deposited to PubMLST, including all human-associated commensal *Neisseria*: *N. benedictiae, N. bergeri, N. blantyrii, N. cinerea, N. elongata, N. lactamica, N. maigaei, N. mucosa, N. oralis, N. polysaccharea, N. subflava, N. uirgultaei* and *N. viridiae*. *N. gonorrhoeae* was included as the focal pathogen in this analysis, however *N. meningitidis* was excluded. We limited the majority of the gonococcal dataset to the last five years (2020-2024) to capture more recent commensal/pathogen transfer events, though included commensals isolates from all years, and added some earlier *N. gonorrhoeae* isolates of interest. Alleles for each gene were aligned using MAFFT v.7 with a UPGMA tree calculated^50^. iTOL was used to visualize and annotate the resultant trees^51^. Multispecies phylogenies were assessed for admixture using the *genealogical sorting index* (*gsi*) with the *genealogicalSorting* R package^52,53^.

## Results

### Phenotypic results of In vitro selection

The four commensal species selected to undergo experimental evolution (AR-0944 [*N. cinerea*], AR-0945 [*N. elongata*], AR-0948 [*N. canis*], and AR-0953 [*N. subflava*]), represented evolutionarily distinct starting points with known MIC values, as outlined in Table 1. Four replicate lineages of each of the four commensal species underwent the aforementioned passaging procedure. In response to ciprofloxacin selection, the average MICs of each species are as follows; *N. cinerea* 7.5 ± 3.4 μg/mL, *N. elongata* 1.38 ± 0.88 μg/mL, *N. canis* 0.5 ± 0.68 μg/mL, and *N. subflava* 12 ± 4.62 μg/mL (Figure 1). The MIC values of 10 evolved lines surpassed the resistance breakpoint of ≥ 1 μg/mL as outlined by CLSI guidelines. The smallest changes in MIC were observed in *N. canis* and *N. elongata*, while all experimental replicates for *N. cinerea* and *N. subflava* surpassed the resistance breakpoint.

**Figure 1.**
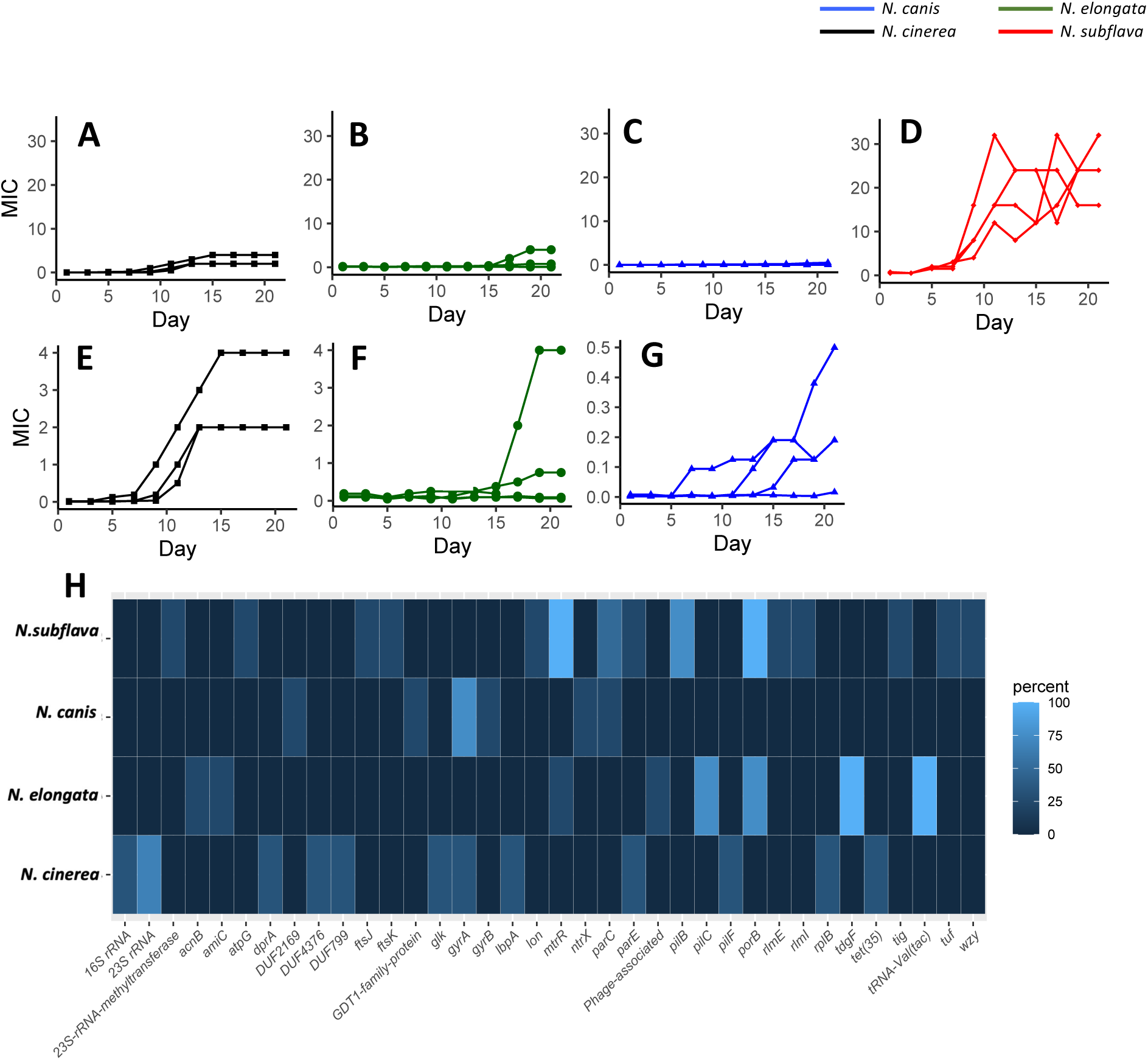
*In vitro* selection of four commensal Neisseria species with ciprofloxacin. In brief, four experimental replicates of each species were passaged for 20 days on selective gradients created with Etest strips. Cells for each passage were selected by sweeping the zone of inhibition (ZOI) and a 1 cm band in the bacterial lawn surrounding the ZOI. MIC trajectories are shown for (A) *N. cinerea*, (B) *N. elongata*, (C) *N. canis*, and (D) *N. subflava*. Expanded scales are shown in panels E-G for *N. cinerea*, *N. elongata*, and *N. canis* respectively. (H) CDs identity of *in vitro* derived mutations for drug-selected lineages. The percentage of mutational hits within a given gene is displayed as a heatmap, with brighter blue coloration indicating more frequent occurrence of a mutation within a CDS in replicate evolved lineages for each species.

### Comparative genomics and nominating derived mutations

Selected lineages were whole genome sequenced (WGS) and aligned to the draft assemblies reported previously^36,37,42^ to nominate derived polymorphisms. Mutations shared with control strains (with no drug selection) were omitted from subsequent analyses. 717 derived mutations were identified across all sequenced strains, averaging 44.81± 33.15 per evolved lineage. The most frequent mutation occurring in *N. cinerea* lineages was located within the *23s rDNA*, the genomic region encoding the 23s rRNA (present in 66% of replicates). Additional derived mutations of note included those in *parE* encoding one of the topoisomerase IV subunits (33%), *gyrA* encoding the DNA gyrase A subunit (33%), and *pilF,* an ATPase that promotes pilus elongation (33%; Figure 1H; see Supplementary Table 1 for complete list). For, *N. elongata* the most frequent mutations were in *tdfG,* a TonB-dependent transporter (100%) and the region encoding the transfer RNA tRNA-Val(tac) (100%), followed by those in regions encoding the major outer membrane porin (*porB;* 75%), a component of the pilus involved in retraction (*pilC;* 75%), the repressor of the *mtr efflux pump* (*mtrR;* 25%), the N-acetylmuramyl-l-alanine amidase (*amiC;* 25%), and aconitate hydratase 2/2 methylisocitrate dehydratase (*acnB;* 25%) (Figure 1H). For *N. canis* the most frequent mutations were in *gyrA* (75%), followed by those in *parC* encoding one of the topoisomerase IV subunits (25%), the a sensor-histidine kinase/response regulator (*ntrX;* 25%), and DNA gyrase subunit B (*gyrB*; 25%) (Figure 1H). For *N. subflava* the most frequent mutations were in *porB* (100%) and *mtrR* (100%); with additional mutations of note in pilB, a cytoplasmic NTPase involved in modulating pilus extension and retraction (75%), *parC* (50%), and *parE* (25%; Figure 1H; see Supplementary Table 1 for complete list).

For particular loci of note, mutations that emerged included: A R320S substitution in topoisomerase IV subunit E and a T91I substitution in DNA gyrase subunit A in *N. cinerea*; a T11N substitution in MtrR in *N. elongata*; substitutions in DNA gyrase subunit A (G89D, S91N, S91R), DNA gyrase subunit B (P739H), and topoisomerase IV subunit C (G85C) for N. canis; and MtrR (*504insA*, *300delA*, T93P, G45S), topoisomerase IV subunit C (A123E, 477 insertion K) topoisomerase IV subunit E (646 insertion R), and reading frame shifts in *porB* in *N. subflava* (Table 2). For specific haplotypes of each evolved lineage see Supplementary Table 2.

**Table 2.** Derived mutations emerging in selected commensal lineages, compared to known ciprofloxacin resistance-contributing loci in *N. gonorrhoeae*.

| Loci contributing to resistance in *N. gonorrhoeae* | Specific mutations in *N. gonorrhoeae* | Derived mutations in commensals |  |  |  |
| --- | --- | --- | --- | --- | --- |
|  |  | *N. canis* | *N. cinerea* | *N. elongata* | *N. subflava* |
| *gyrA* | S91F, D95A/N/G | G89D*, S91N*, S91R* | T91I |  |  |
| *gyrB* | D429N | P739H* |  |  |  |
| *mtrR* | A39T, G45D, Y105H |  |  | T11N* | 504ins-a<sup>2</sup>, 300del-a<sup>2</sup>, T93P, G45S |
| *parC* | D86N, S87R/N, E91K | G85C* |  |  | A123E, 477insK |
| *parE* | G410V |  | R320S |  | 646insR |
| *porB* | E127G |  |  |  | KO-indels<sup>2</sup> |

\*Strains evolving these mutations did not cross the reduced susceptibility breakpoint of $\geq 1$ $\mu\text{g/mL}$ .
<sup>2</sup>-Amino acid residue numbering scheme is based on alignment to the gonococcal reference genome (NZ\_AP023069.1).
<sup>2</sup> Putative DNA-level insertions or deletions of nucleotides.

### Testing for zoliflodacin cross-resistance in evolved cell lines

We tested evolved lineages for the emergence of cross-resistance to zoliflodacin after ciprofloxacin selection. Though no strains displayed large changes in zoliflodacin MICs compared to ancestral lineages (Table 3): 1 *N. elongata* replicate had an elevated MIC (from 1 to 4 μg/mL), 1 *N. canis* replicate had an elevated MIC (from 1 to 4 μg/mL), and all 4 *N. subflava* lineages had elevated MIC (from 2 to 4 μg/mL). For the *N. elongata* isolate with an elevated zoliflodacin MIC, it was the only lineage to inherit a MtrR T11N substitution post-selection. For the *N. canis* isolate with an elevated zoliflodacin MIC, it was the only to inherit GyrA G89D and ParC G85C mutations. Finally, for the *N. subflava* lineages, all inherited a MtrR mutation (300delA, T93P, or G45S); coupled with *porB* indels which altered the reading frames; with 3 out of the 4 lineages also inheriting either a K insertion at position 477 in ParC, ParC A123E, or a R insertion at position 646 in ParE.

**Table 3.** Characterization of zoliflodacin MICs for ciprofloxacin-selected lineages.

|  | Zoliflodacin MICs (µg/mL) |  |  |  |  |
| --- | --- | --- | --- | --- | --- |
| Species | Ancestral | Replicate 1 | Replicate 2 | Replicate 3 | Replicate 4 |
| AR-0944 | 2 | 2 | 1 | 1 | 0.25 |
| AR-0945 | 1 | 1 | 1 | 1 | 4 |
| AR-0948 | 1 | 4 | 1 | 1 | 1 |
| AR-0953 | 2 | 4 | 4 | 4 | 4 |

**\*DNG=Single colony picks did not grow.**

### Population analysis of derived mutations and HGT in natural Neisseria populations

We searched for evidence of *in vitro* evolved mutations and gene-level recombination tracts in natural *Neisseria* populations using *Neisseria* isolates deposited to the PubMLST database. For the *gyrA* locus, we observed an *in vitro* selected mutation at position 89 (G89D); however, we found no evidence of amino acid substitutions at this position in natural populations of *Neisseria,* with all isolates (n=2231) harboring a glycine at that position. At position 91 however, we find a diversity of alleles encoding different amino acids in natural *Neisseria* populations with: 1 isolate harboring an alanine (A), 489 with a phenylalanine (F), 70 with an isoleucine (I), 1 with a leucine (L), 531 with a serine (S) 1128 with a threonine (T), 10 with a valine (V), and 1 with a tyrosine (T) (Figure 2A). Of these mutations, 1 was uncovered in our experimental evolution experiments 91I and was associated with elevated ciprofloxacin MICs. 91I in natural populations of *Neisseria* was found in *N. lactamica*, *N. subflava*, and *N. polysaccharea* (Figure 2A). We find no evidence of 91F in commensals which is associated with ciprofloxacin resistance in gonococci (Figure 2A). Though we did not find any evolved mutations at position 95 in GyrA in our *in vitro* selection, we characterized the amino acid diversity across natural populations as D95A/N/G substitutions increase ciprofloxacin MICs in gonococci. Here we find, 369 isolates harboring an alanine (A), 1708 with a aspartic acid (D), 118 with a glycine (G), 1 with a histidine (H), 25 with an asparagine (N), and 10 with a tyrosine (Y) (Figure 2A). Reduced susceptibility-associated amino acids were found in *N. lactamica*, *N. subflava*, *N. polysaccharea* and *N. bergeri*. We find no evidence of admixture between commensals and gonococci at the locus-level for *gyrA* however in this analysis, with gonococci belonging to a monophyletic clade (*gsi* = 1).

**Figure 2.**
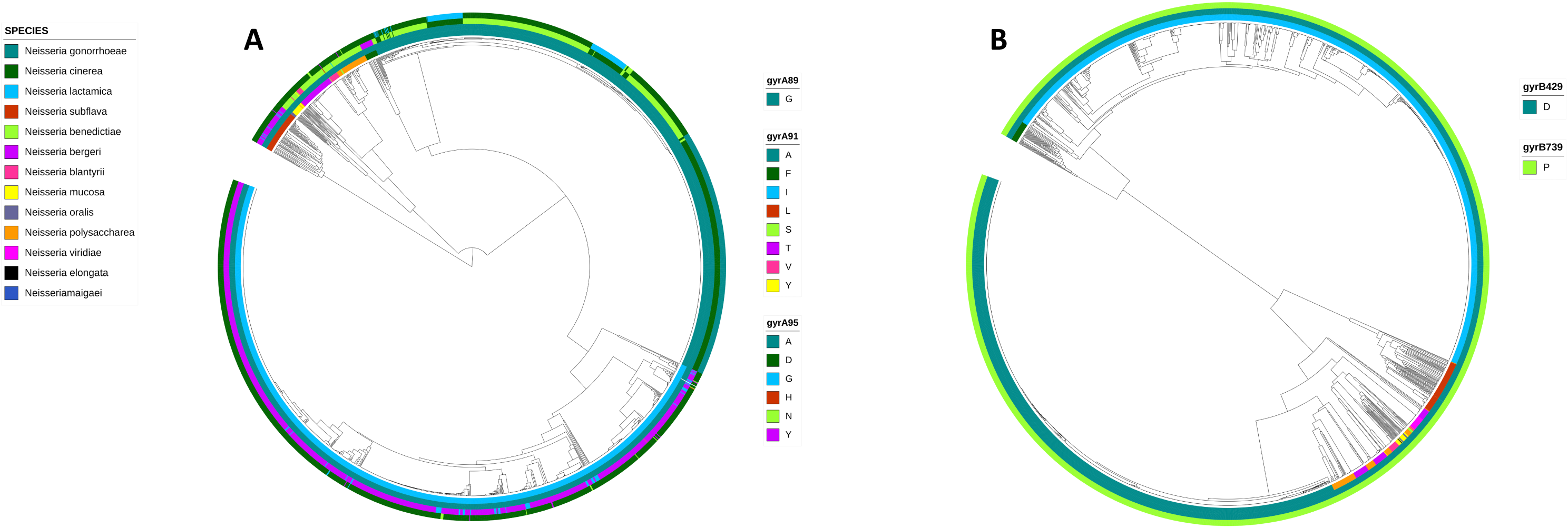
(A) UPGMA phylogeny of *gyrA* of 2235 PubMLST *Neisseria* isolates. The ciprofloxacin reduced susceptibility-associated mutation GyrA 91I circulates within commensal *Neisseria* populations, however has not been transferred to *Neisseria gonorrhoeae*. The coloring in the inner ring represents the *Neisseria* species, followed by amino acid identity at GyrA positions 89, 91, and 95 respectively. The derived 91I mutation was found in *N. lactamica*, *N. subflava*, and *N. polysaccharea* (n=70 or ∼ 3% of isolates). (B) UPGMA phylogeny of the *gyrB* locus for 2302 PubMLST *Neisseria* isolates. GyrB does not show variation in the zoliflodacin reduced susceptibility mutation (D429V) or derived P739H mutation in commensal populations. The coloring in the inner ring represents the *Neisseria* species, followed by amino acid identity at GyrB positions 429 and 739 respectively.

In our selection experiment we uncovered a GyrB P739H substitution emerge post-selection in *N. canis*. In natural *Neisseria* populations however, we did not find any evidence of this mutation, with all isolations (n=2301) homozygous for a proline (P) at this position (Figure 2B). Additionally, we searched for the D429N substitution that is associated with zoliflodacin in gonococci^31^. All isolated harbored an aspartic acid (D) at this position however, with no variation observed (Figure 2B). Finally, we find no evidence of commensal to pathogen allelic exchange at the *gyrB* locus level (gonococcal *gsi* = 1).

At *parC*, a mutation encoding a G85C substitution emerged in *N. canis,* and A123E and 477 insertion K substitutions emerged in *N. subflava* after selection. In natural *Neisseria* populations we find no variation at position 126 or 477. We next investigated the positions 85, 86, 87, and 91; as we found post-selection variation after experimental evolution at position 85, and all three of the later residues are associated with ciprofloxacin resistance in gonococci. Here we find: 2231 isolates with a glycine (G), 23 with a serine (S), 1 with a cysteine (C), and one with an aspartic acid (D) at position 85; 2181 isolates harboring an aspartic acid (D), and 75 with a asparagine (N) at residue 86; 1911 with a serine (S), 318 with an arginine (R), 21 with an asparagine (N), and 6 with an isoleucine (I) at position 87; and 2208 with a glutamic acid (E), 33 with a glycine (G), 3 with a lysine (K), and 12 with a glutamine (Q) at position 91 (Figure 3A). For *parC*, we also identify a putative gene-level HGT event, with a clade of three gonococcal isolates from the Netherlands collected in 2021 harboring more *N. polysaccharea* or *N. lactamica*-like alleles (Figure 3A). All three isolates had haplotypes of 86D 87S 91E. The 86N, 87R/N, and 91K mutations associated with ciprofloxacin resistance in gonococci were limited to the gonococcal clade and were not present in the commensal *Neisseria* (Figure 3A). We also identify a single gonococcal isolate collected in China with the ParC G85C substitution that emerged in *N. canis* post-selection (Figure 3A).

**Figure 3.**
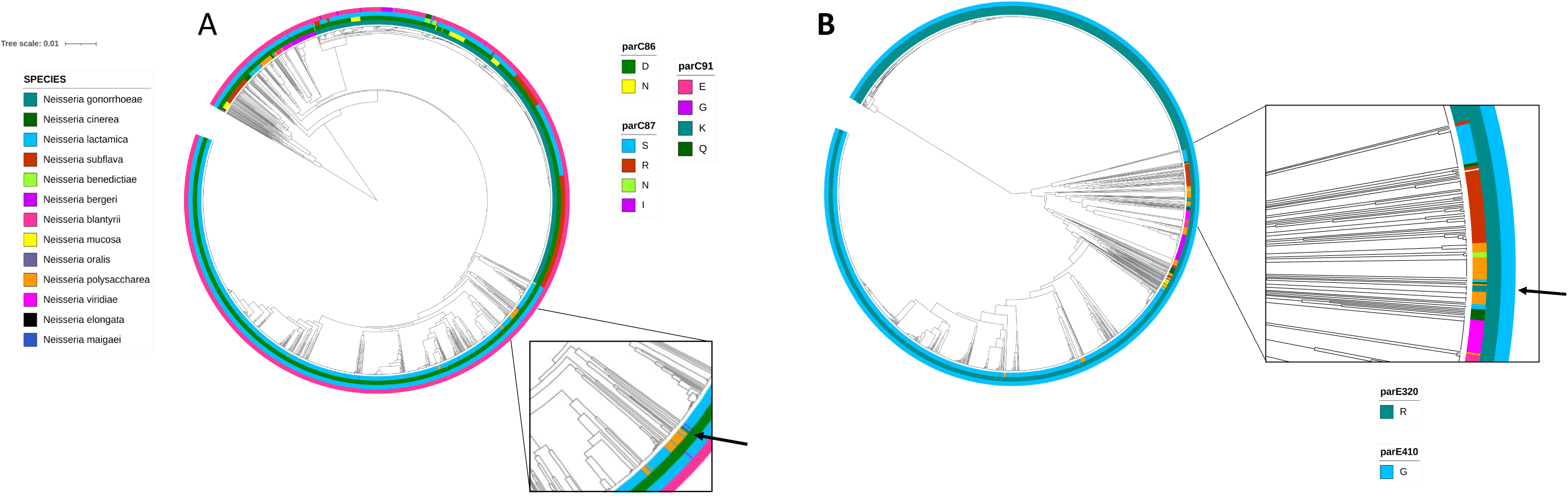
(A) UPGMA phylogeny of the *parC* locus for 2256 PubMLST *Neisseria* isolates. ParC 86N, 87R/N, and 91K mutations associated with ciprofloxacin resistance in gonococci were limited to the gonococcal clade and were not present in the commensal *Neisseria*; however, a single gonococcal isolate inherited the experimentally-derived ParC G85C substitution that emerged in *N. canis* post-selection. The coloring in the inner ring represents the *Neisseria* species, followed by amino acid identity at ParC positions 86, 87, and 91 and respectively. The subpanel highlights 3 gonococcal isolates with commensal-like *parC* alleles. (B) UPGMA phylogeny of the *parE* locus for 2287 PubMLST *Neisseria* isolates. The coloring in the inner ring represents the *Neisseria* species, followed by amino acid identity at ParE positions 320 and 410 respectively. The subpanel highlights 4 gonococcal isolates with commensal-like *parE* alleles.

For ParE, substitutions emerging after ciprofloxacin selection included R320S in *N. cinerea* and 646 R insertion in *N. subflava*. We found no evidence of variation at position 320 in natural Neisseria populations, with all isolates (n=2286) harboring a R. We also looked for evidence of variation at position 410 which has been associated with x resistance in gonococci, however we also find no variation at this site in natural *Neisseria* populations. We do however find HGT of commensal Neisseria *parE* alleles into a cluster of four gonococcal isolates from Scotland isolated in 2023-24 (Figure 3B).

## Discussion

Cross-species transfer of alleles harboring ciprofloxacin resistance-associated mutations has been observed in natural *Neisseria* populations^5,48^, where commensals serve as the reservoir for pathogen evolution. Thus, characterizing the mechanisms that impart ciprofloxacin resistance in *Neisseria* commensals using *in vitro* selection may highlight important mutations to consider for gonococcal surveillance programs and may also illuminate novel resistance-conferring mutations that can arise in commensal genomic backgrounds. Previous work using *in vitro* evolution has pointed to conserved mechanisms of resistance across the genus for azithromycin and penicillin-based selection^36,37,42^, here we expand our enquiry to ciprofloxacin for several *Neissiera* commensals, and interrogate both the identity and diversity of paths to resistance across different genomic backgrounds.

Ciprofloxacin, a quinolone-class antibiotic, targets the bacterial type II topoisomerases DNA gyrase and topoisomerase IV, and disrupts DNA synthesis; therefore it is no surprise that in gonococci, reduced susceptibility is often mediated by mutations in the primary and secondary drug targets, GyrA and ParC respectively^12,20,24,25^. In particular, mutations falling within the quinolone resistance-determining regions (QRDRs) of GyrA and ParC (codons 91-95 of GyrA and 86-91 of ParC) have been shown to impact ciprofloxacin MICs, with the substitutions S91F and D95A/N/G in GyrA and D86N, S87R/N, and E91K ParC strongly associated with elevated MICs in natural gonococcal populations^12,31^. Here in *Neisseria* commensals, we find amino acid-altering mutations in these two loci: G89D, S91N, S91R, and T91I in GyrA; and G85C, A123E, and a K insertion at position 477 in ParC (Table 2); the majority of which fall within the aforementioned QRDRs. Of these mutations T91I has been found in quinolone resistant meningococci with gyrA alleles that had been inherited from commensal *Neisseria* donors (*N. lactamica*, *N. cinerea*, and *N. subflava*), and is associated with elevated ciprofloxacin MICs in conjunction with ParC mutations^5^. Here, we find this 91I mutation in natural commensal populations of *Neisseria* (*N. lactamica*, *N. subflava*, and *N. polysaccharea*; n=70 or ∼ 3% of isolates) suggesting its availability for HGT and resistance transfer, however we do not see this mutation in contemporary gonococci (Figure 2A). Finally, we also find the ParC G85C substitution which emerged post-selection in a single gonococcal isolate. The isolate was collected in China (ciprofloxacin MIC of 2 μg/mL) and had resistance-associated mutations at GyrA 91F and 95A, however had wildtype alleles at ParC at other resistance associated sites (86D, 87S, 91E). Thus, we believe the ParC G85C substitution is acting together with GyrA 91F and 95A to produced elevated ciprofloxacin MICs in this strain.

We also find substitutions that emerged in the other subunits of DNA gyrase and topoisomerase IV: GyrB P739H, ParE R320S, and a R insertion at position 646 of ParE. There was no variation at these sites in the natural commensal or gonococcal populations included within this analysis (Figure 2B and Figure 3B). The GyrB P739H has been associated with intermediate ciprofloxacin resistance (MICs: 0.032 to 0.125 mg/L) in gonococci in the laboratory, and has also been observed in two commensal species that were not included in our population analysis (*N. brasiliensis* (PATRIC ID-2666100.4) and an unidentified *Neisseria* isolate (PubMLST ID-94179))^30^. Low rates of carriage likely suggest some fitness cost to maintenance of GyrB P739H, though follow up experiments will be necessary to understand the impact on fitness for all uncovered mutations.

Investigation of the haplotypes of each evolved lineage can help us to understand which mutations may be contributing to observed ciprofloxacin resistance. For *N. cinerea*, though we were unable to identify the mutations contributing to resistance in 3 strains, the fourth replicate inherited both GyrA T91I and ParE R320S (MIC = 6 µg/ml; Supplemental Table 2). We suspect that both of these mutations contribute together to the high ciprofloxacin MICs in this strain, as in meningococci the GyrA T91I must be inherited together with topoisomerase IV mutations (ParC D86N, S87R/I, E91G) to produce elevated ciprofloxacin MICs^5^. In *N. canis*, a similar pattern exists where GyrA or GyrB substitutions only marginal increase ciprofloxacin MICs compared to the ancestral strain, however a strain inheriting both GyrA G89D and ParC G85C had a MIC above the breakpoint (MIC = 1.5 µg/ml; Supplemental Table 2). For *N. subflava*, the largest contributors to elevated MICs are less clear, with all strains inherited 2-3 likely contributors; however, we speculate the combination of reduced drug influx from porin knockouts and increased drug efflux from *mtrR* mutations are likely causing the high MICs observed in these strains (≥ 8 µg/ml) as they all lack the combination of GyrA/B and ParC/E mutations typically required to elevated ciprofloxacin MICs in Neisseria.

Emergence of ciprofloxacin-selected mutations in type II topoisomerases may be concerning given the current development of zoliflodacin as a new treatment for gonorrhea. Zoliflodacin has a novel target (GyrB) and mode of action compared to ciprofloxacin^41^, however we questioned if zoliflodacin cross-resistance could emerge post-ciprofloxacin selection given that mutations emerged in GyrB. Prior mutations associated with elevated zoliflodacin MICs include GyrB D429V, GyrB S467N, and GyrB K450^31,54^ which did not emerge here (Table 2), however we did recover a derived GyrB P739H mutation. The *N. cinerea* strain that inherited GyrB P739H did not have an elevated zoliflodacin MIC compared to the ancestral strain; yet interestingly other selected strains did (Table 3). These strains included: A *N. elongata* strain (MIC increase from 1 to 4 µg/ml) inheriting a MtrR T11N substitution, a *N. canis* strain (MIC increase from 1 to 4 µg/ml) inheriting GyrA G89D and ParC G85C substitutions, and all four *N. subflava* strains (MIC increase from 2 to 4 µg/ml) which had a combination of influx, efflux, and singleton ParC or ParE mutations (Table 3 and Supplementary Table 2). Upon investigating the haplotypes of these strains ParC and ParE mutations appeared to have a limited impact on zoliflodacin MICs in *N. subflava* as the strain without them also had a zoliflodacin MIC of 4 µg/ml (Table 4 and Supplementary Table 2); thus, we speculate the majority of cross-resistance observed is emerging through generalized mechanisms (i.e., drug influx or efflux), aside from the single *N. canis* isolate with the GyrA G89D and ParC G85C substitutions. As GyrB variation in natural *Neisseria* populations is rare (^54^ and Figure 2B), future studies may consider testing the impact of naturally occurring efflux pump and porin mutations on zoliflodacin MIC when assessing the current and future potential for resistance emergence rather than focusing on variation in the drug target alone.

In conclusion, this work again highlights the potential of commensal *Neisseria* in serving as reservoirs of resistance for their pathogenic relatives. We find evidence of recent transfer of *parC* and *parE* genes from commensals to pathogens through the observation of admixed phylogenies (Figure 3A and 3B). We also characterize mutations associated with ciprofloxacin reduced susceptibility in commensals after selection, and find both previously described and novel mutations emerging in the drug targets, the Mtr efflux pump system, and the major outer membrane porin. Of these mutations we identify GyrA 91I as a substitution circulating within commensal *Neisseria* populations, confirming prior results^5^; and a ParC G85C present in a single ciprofloxacin-resistant gonococcal isolate. Finally, we observe cross-resistance to zoliflodacin emerge, and speculate drug influx and/or efflux mutations are the likely underlying contributors.

## Statement

Whole genome sequencing libraries can be accessed through the NCBI’s Sequence Read Archive (SRA) with the accession numbers: SAMN41424178-SAMN41424193.

## Acknowledgements

Research by C.B.W. is supported by the College of Science and Thomas H. Gosnell School of Life Sciences at RIT and the National Institute of General Medical Sciences of the National Institutes of Health under Award Number R15AI174182. The content is solely the responsibility of the authors and does not necessarily represent the official views of the National Institutes of Health.

